# Automated identification of cell-type–specific genes and alternative promoters

**DOI:** 10.1101/2021.12.01.470587

**Authors:** Mickaёl Mendez, Jayson Harshbarger, Michael M. Hoffman

## Abstract

**Background:** Identifying key transcriptional features, such as genes or transcripts, involved in cellular differentiation remains a challenging problem. Current methods for identifying key transcriptional features predominantly rely on pairwise comparisons among different cell types. These methods also identify long lists of differentially expressed transcriptional features. Combining the results from many such pairwise comparisons to find the transcriptional features specific only to one cell type is not straightforward. Thus, one must have a principled method for amalgamating pairwise cell type comparisons that makes full use of prior knowledge about the developmental relationships between cell types.

**Method:** We developed Cell Lineage Analysis (CLA), a computational method which identifies transcriptional features with expression patterns that discriminate cell types, incorporating Cell Ontology knowledge on the relationship between different cell types. CLA uses random forest classification with a stratified bootstrap to increase the accuracy of binary classifiers when each cell type have a different number of samples. Regularized random forest results in a classifier that selects few but important transcriptional features. For each cell type pair, CLA runs multiple instances of regularized random forest and reports the transcriptional features consistently selected. CLA not only discriminates individual cell types but can also discriminate lineages of cell types related in the developmental hierarchy.

**Results:** We applied CLA to Functional Annotation of the Mammalian Genome 5 (FANTOM5) data and identified discriminative transcription factor and long non-coding RNA(lncRNA) genes for 71 human cell types.With capped analysis of gene expression (CAGE) data, CLA identified individual cell-type–specific alternative promoters for cell surface markers. Compared to random forest with a standard bootstrap approach, CLA’s stratified bootstrap approach improved the accuracy of gene expression classification models for more than 95% of 2060 cell type pairs examined. Applied on 10X Genomics single-cell RNA-seq data for CD14^+^ monocytes and FCGR3A^+^ monocytes, CLA selected only 13 discriminative genes. These genes included the top 9 out of 370 significantly differentially expressed genes obtained from conventional differential expression analysis methods.

**Discussion:** Our CLA method combines tools to simplify the interpretation of transcriptome datasets from many cell types. It automates the identification of the most differentially expressed genes for each cell type pairs CLA’s lineage score allows easy identification of the best transcriptional markers for each cell type and lineage in both bulk and single-cell transcriptomic data.

**Availability:** CLA is available at https://cla.hoffmanlab.org. We deposited the version of the CLA source with which we ran our experiments at https://doi.org/10.5281/zenodo.3630670. We deposited other analysis code and results at https://doi.org/10.5281/zenodo.5735636.

## 1 Introduction

Discovering cell-type–specific transcriptional features is important for identifying oncogenes^1^, finding disease-associated genes^2–4^, and determining drivers of cell differentiation^5^. Current methods allow identifying the transcriptional features differentially expressed between a pair of cell types^6–8^. These differential expression analysis methods, however, often identify a long list of significant transcriptional features of variable relevance. Ideally, one would want a method to focus on the most important transcriptional features that can discriminate among cell types. The availability of large numbers of expression datasets represents an opportunity to identify cell-type–specific transcriptional features by aggregating the results from multiple cell type pairs.

To simplify the analysis of multiple cell type pairs we have developed the Cell Lineage Analysis (CLA) framework, which uses the regularized random forest^9^ machine learning classifier. CLA indicates how often a particular transcriptional feature discriminates one cell type in multiple cell type pairs. CLA also uses information from the Cell Ontology^10^ about the relationships between cell types to find the transcriptional features that discriminate lineages.

The Cell Ontology describes hierarchical relationships between most cell types using relations such as *is_a* (Figure 1). The *is_a* relation implies inheritance between cell types such that the children inherit the properties of their ancestors. By using the *is_a* relationships, we define a cell type’s *lineage* as that cell type and all its descendants.

**Figure 1:**
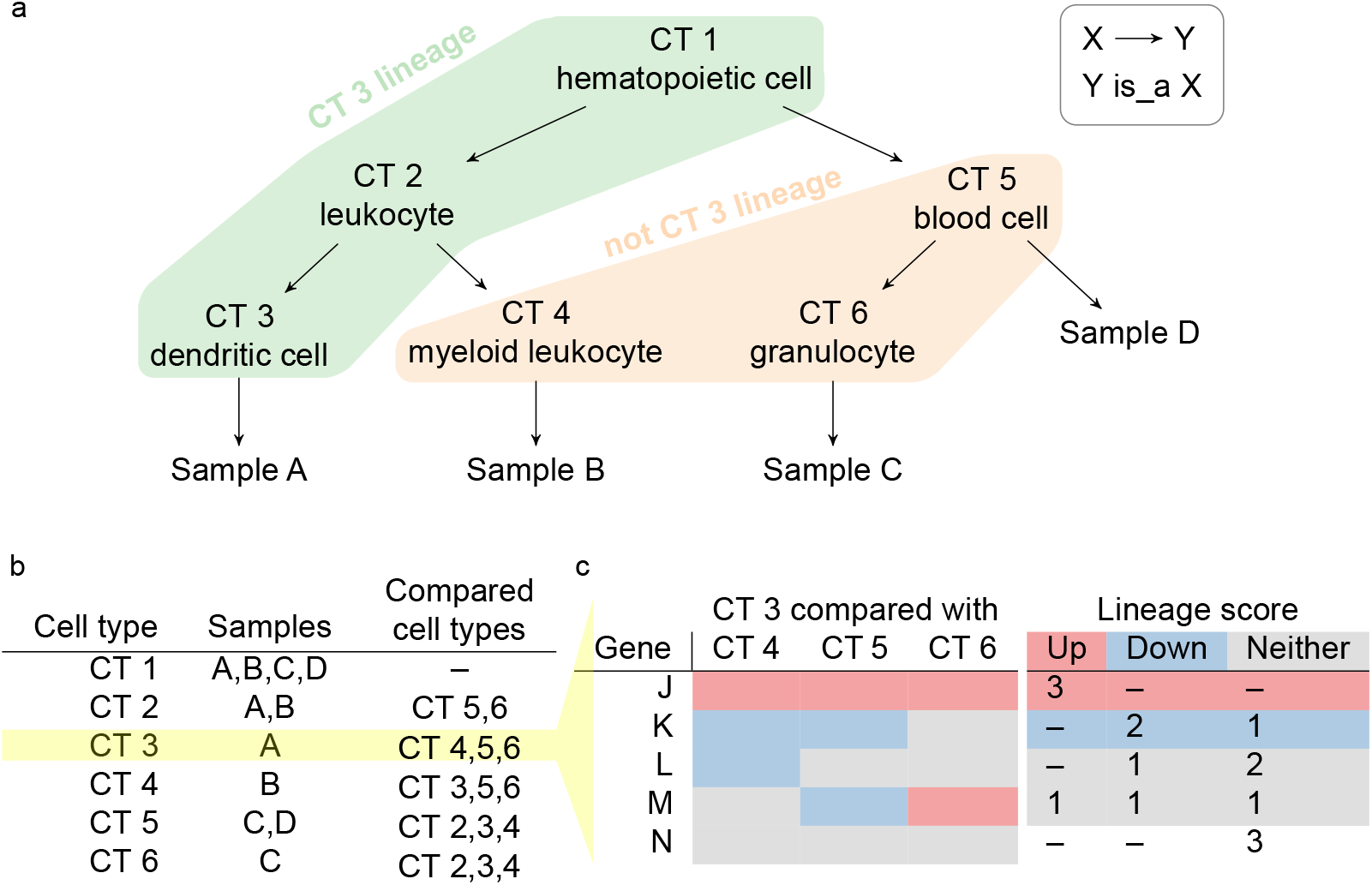
Summary workflow to compute lineage scores. **(a)** Toy example of the Cell Ontology graph as a tree for hematopoietic cell lineage. Green: Lineage of dendritic cell (CT 3), including its 2 ancestors leukocyte (CT 2) and hematopoietic cell (CT 1); orange: 3 cell types not in the CT 3 lineage, including siblings and cousins of dendritic cell. Leafs: 4 physical hematopoietic cell samples A–D. Arrows: *is_a* relation between either two cell types or one cell type and a sample of that type. **(b)** The 6 cell types, and for each cell type, the associated samples and the other cell types CLA would compare against—those cell types neither ancestors or descendants of the first cell type. Yellow: an illustrative case of dendritic cells (CT 3). **(c)** 5 genes J–N discriminating dendritic cells (CT 3) from myeloid leukocytes (CT 4), blood cells (CT 5), and granulocytes (CT 6). (*Left*) Expression pattern of the selected genes. Red: higher expression in dendritic cell samples (Up);blue: lower expression in dendritic cell samples (Down); grey: not selected (Neither). (*Right*) Lineage score indicating the number of times that each gene had higher (red), lower (blue), or non-discriminative expression (grey) in dendritic cells in most comparisons. White row: gene has non-discriminative expression in dendritic cells in all comparisons.

CLA uses the FANTOM5 (FF) sample ontology^11^ which assigned cell line and primary cell samples from Functional Annotation of the Mammalian Genome 5 (FANTOM5) to matching cell types in the Cell Ontology.

For cell types within lineages, CLA uses the *is_a* relationship such that each cell type inherits the samples from its descendants. Thus, if CLA identifies a transcriptional feature that discriminates both a cell type and its parent cell type against other cell types, we describe that feature as discriminating a lineage.

To identify the transcriptional features that discriminate between a pair of cell types, CLA uses regularized random forest^9^, a machine learning algorithm that builds an ensemble of decision trees that perform classification together. Similar to random forest^12,13^, regularized random forest creates decision points that use transcriptional feature expression values to split samples into two groups. At each decision point, random forest selects one transcriptional feature, out of a random subset of features, which has expression values that discriminate the cell types the most. After creating all the decision points, one can estimate the discriminative power of the selected features.

To remove the least discriminative transcriptional features, and thus reduce the number of features selected by random forest, regularized random forest adds a regularization step that penalizes the selection of new features during the construction of decision trees. Regularized random forest selects fewer transcriptional features when compared to the non-regularized version. Therefore, regularized random forest increases the chances of selecting the most discriminative transcriptional features.

In addition to standard Cell Ontology information, CLA requires two inputs: (1) a table of expression values for transcriptional features in different samples and (2) a list of associations between samples and cell types. As individual samples can be annotated with multiple cell types from the same lineage, we excluded comparisons that would involve the same sample in both cell types.

Here, we apply CLA to 751 FANTOM5 human primary cell and cell line samples grouped into 71 Cell Ontology cell types. We show that CLA identifies cell-type–specific genes from a transcription factor dataset and a long noncoding RNA (lncRNA) dataset. We also show that it identifies alternative promoters from a cluster of differentiation (CD) dataset.

Our CLA software automates the exploratory analysis of cell type pairs. CLA also identifies the most differentially expressed genes from peripheral blood mononuclear single-cell RNA-seq data. We provide the data via CLA’s user-friendly web interface (https://cla.hoffmanlab.org). We implemented CLA using Python 3^14^ and scikit-learn^15^ (https://github.com/hoffmangroup/cla).

## 2 Results

### 2.1 CLA discriminates cell types with higher accuracy than the unstratified approach

Due to the hierarchical structure of the Cell Ontology, each cell type may associate with different number of samples. For example, parent cell types, such as hematopoietic cell ([CL:0000988](https://www.ebi.ac.uk/ols/ontologies/cl/terms?iri=http%3A%2F%2Fpurl.obolibrary.org%2Fobo%2FCL_0000988); *n* = 182), have more samples than descendant cell types such as melanocyte ([CL:0000148](https://www.ebi.ac.uk/ols/ontologies/cl/terms?iri=http%3A%2F%2Fpurl.obolibrary.org%2Fobo%2FCL_0000148); *n* = 12). This leads to cell type comparisons with class imbalance, a problem known to impede the performance of decision trees^16^.

We designed CLA so that it would work well in conditions of class imbalance. To reduce biases from class imbalance, we used a stratified sampling approach for random forest^17^. The stratified sampling approach starts growing each tree by selecting randomly, and without replacement, the same number of samples for each cell type pair. We set the number of samples selected per tree to the number of samples from the cell type with fewer samples. To estimate the accuracy of each random forest, we used the unselected samples, also known as out-of-bag samples, and computed the out-of-bag score.

We applied CLA to 751 FANTOM5 human samples from 71 Cell Ontology cell types. Of the 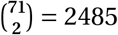 possible cell type pairs, we only selected the 2060 pairs that share zero samples (Figure 1). To evaluate the effect of class imbalance correction with the stratified sampling approach, we compared the accuracies obtained in the 2060 cell type pairs with and without stratified bootstrap. More random forests had higher accuracy with stratified approach than without (Figure 2a).

**Figure 2:**
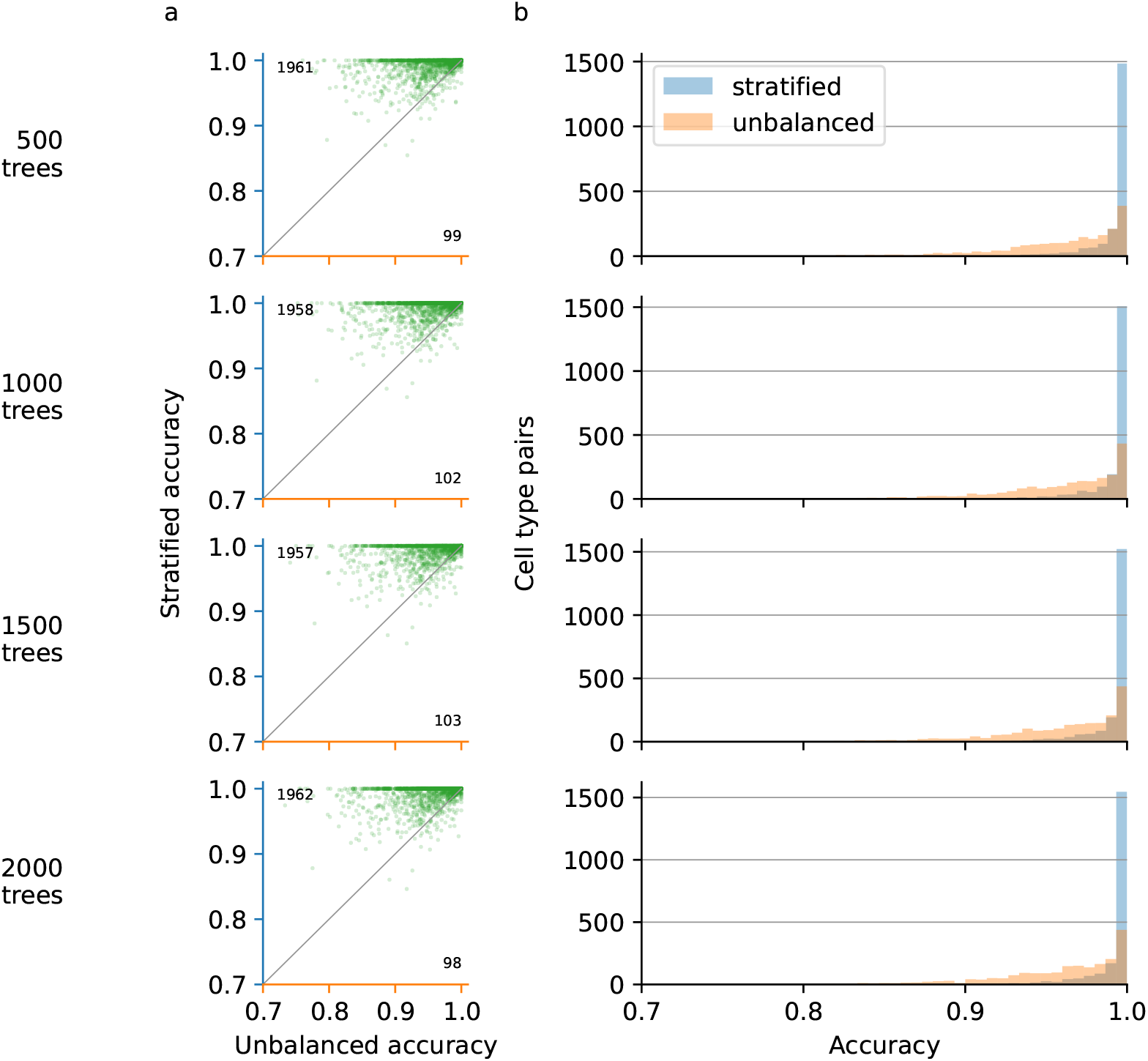
Number of correctly classified samples out of all samples (accuracy) obtained with random forest on 2060 cell type pairs and expression of 1639 transcription factors. Rows: number of trees used. **(a)** Accuracy of each cell type pair with stratified bootstrap against accuracy without stratified bootstrap. Numbers indicate how many cell type pairs have greater accuracy when using stratified bootstrap (top left) or without stratified bootstrap (bottom right). **(b)** Histogram of accuracies for each cell type pair with stratified bootstrap (blue) and without stratified bootstrap (orange).

With 500, 1000, 1500, and 2000 trees, the average accuracy of the 2060 classifiers ranged between 99.30% and 99.31%. Thus, increasing the number of trees for each random forest had no noticeable effect on the average accuracy across all classifiers.

Out of 2060 random forest binary classifiers with 500 trees, 1961 had a higher accuracy with stratified bootstrap than with unbalanced bootstrap (Figure 2b). Using 2000 trees and the stratified bootstrap increased the accuracy of 1962 classifiers. Increasing the number of trees from 500 to 2000 only increased the accuracy of 1 additional cell type pair. In every case, stratified bootstrap increased the accuracy of more than 95% of the classifiers.

### 2.2 Regularized random forest identifies transcriptional features that discriminate pairs of cell types

Regularized random forest may select less discriminative transcriptional features in the first trees and the best transcriptional features in later trees. To avoid reporting of less discriminative features, we ran regularized random forest 10 times for each comparison between a pair of cell types. We defined a robustness score that indicates in how many of these 10 times regularized random forest selected each transcriptional feature. Robustness of 0/10 indicates that regularized random forest never selected the transcriptional feature, and robustness of 10/10 indicates that regularized random forest selected the transcriptional feature every time (Figure 3).

**Figure 3:**
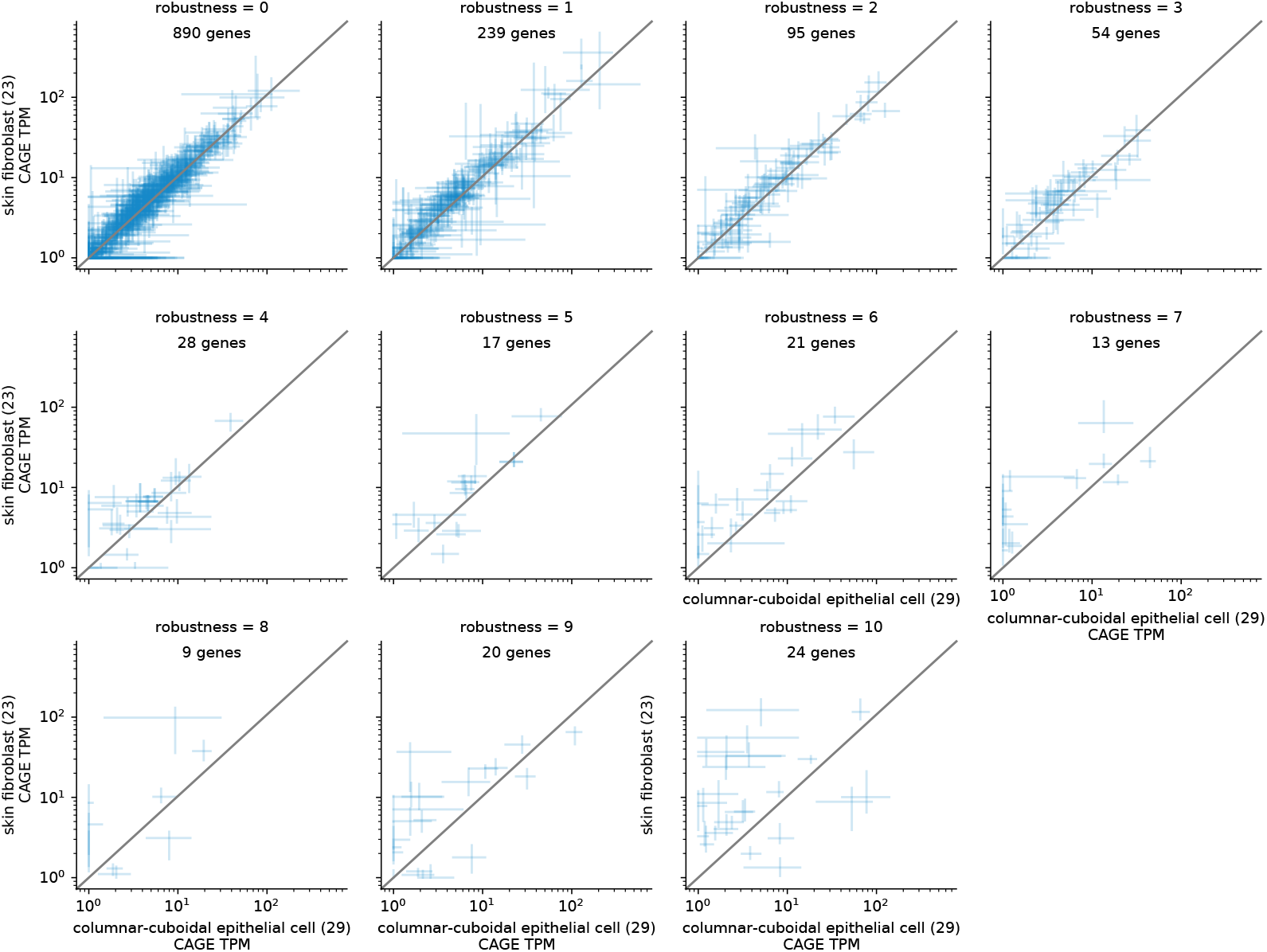
capped analysis of gene expression (CAGE) expression of genes selected in the comparison between 29 columnar-cuboidal epithelial cell samples (CL:0000075) samples and 23 skin fibroblast samples (CL:0002620). Robustness: number of times that CLA selects a gene out of 10 runs. Error bars indicate the first and third quartiles of expression in tag per million (TPM) with intersection at median. Diagonal grey line: *y* = *x* line.

Transcriptional features selected in at least one instance of regularized random forest (robustness > 0) had higher absolute difference of expression in one of the two cell types (mean: 5.67 TPM) than transcriptional features selected in zero instances of regularized random forest (robustness = 0; mean: 1.05 TPM). As we increased the robustness threshold, fewer genes had their first or third quartile expression value near the *y* = *x* line. This indicated that the genes with the highest robustness had distinct expression patterns in the cell type pair. For subsequent analyses, we sought to minimize the number of genes per comparison, focusing only on the most discriminative genes. Thus, below we use only genes with the highest robustness (10/10).

### 2.3 CLA finds lineage-specific transcription factors

To find transcription factors that can alone discriminate different particular cell lineages, we examined the transcription factors CLA selected for individual cell types. We anticipated that cell-type-specific transcription factors would have higher expression in one cell type or lineage than in every other cell type. Thus, we expected that CLA would repeatedly select highly specific transcription factors in comparisons involving the highest number of cell types.

CLA selected the *SPI1* hematopoietic transcription factor^18,19^ in 534/2060 cell type pairs. These 534 pairs involve 70 of the 71 cell types (Figure 4). CLA selected only 11 other transcription factors that provided discrimination for comparisons involving this many cell types. On average, transcription factors discriminate comparisons involving only 31 cell types. This suggests that *SPI1* expression either discriminates 1 cell type against every other cell type, or that it discriminates 1 lineage against many other cell types.

**Figure 4:**
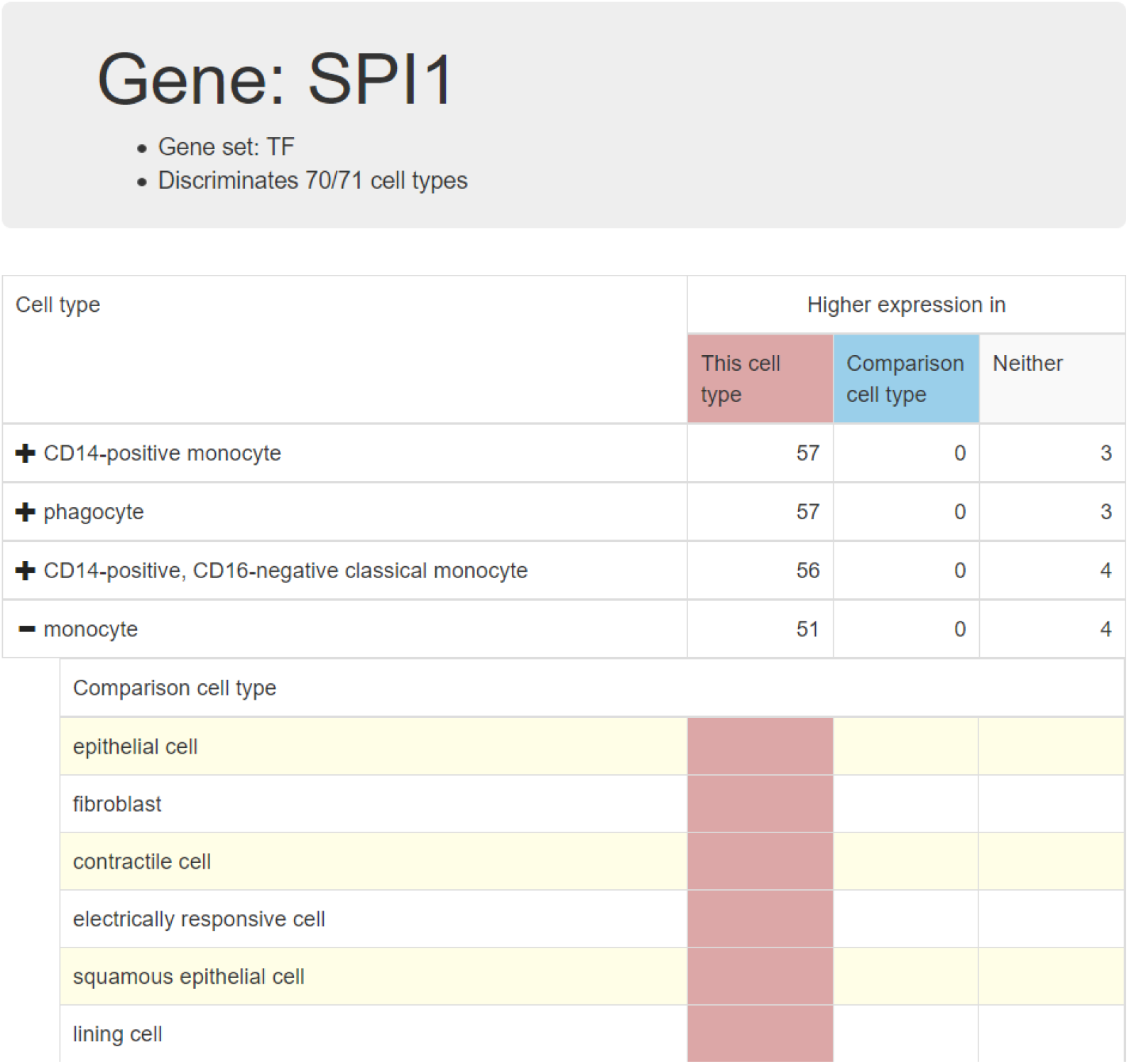
Screenshot of CLA web interface showing *SPI1* lineage scores in the transcription factor dataset. *(Top)* Header, showing gene symbol, gene set, and the fraction of cell types that the gene discriminates. *(Middle)* Lineage scores for each cell type. Red: Number of times that the gene has higher expression in this cell type. Blue: Number of times that the gene has higher expression in the comparison cell type. Grey: Number of times that the gene has no discriminative power. *(Bottom)* Cell types compared with the selected cell type, monocyte. Color indicates if the gene has higher expression in this cell type (red), higher expression in the comparison cell type (blue), or no discriminative power (grey). Every other row has a yellow background as a visual aid.

Out of the 534 cell type pairs, 443 contained one of the 18 cell types from the hematopoietic lineage and 33 contained two cell types from the hematopoietic lineage. In addition, 50 cell type pairs contained phagocytes and a cell type outside of the hematopoietic lineage. While phagocyte ([CL:0000234](https://www.ebi.ac.uk/ols/ontologies/cl/terms?iri=http%3A%2F%2Fpurl.obolibrary.org%2Fobo%2FCL_0000234)) does not have hematopoietic cell ([CL:0000988](https://www.ebi.ac.uk/ols/ontologies/cl/terms?iri=http%3A%2F%2Fpurl.obolibrary.org%2Fobo%2FCL_0000988)) as an ancestor, it does have the hematopoietic cell type CD14-positive, CD16-negative classical monocytes ([CL:0002057](https://www.ebi.ac.uk/ols/ontologies/cl/terms?iri=http%3A%2F%2Fpurl.obolibrary.org%2Fobo%2FCL_0002057)) as a child. Through this child, the phagocyte cell type inherits hematopoietic samples. Similar to phagocyte, we found 5 cell type pairs containing motile cell ([CL:0000219](https://www.ebi.ac.uk/ols/ontologies/cl/terms?iri=http%3A%2F%2Fpurl.obolibrary.org%2Fobo%2FCL_0000219)), a parent of phagocyte, and one cell type pair contained nucleate cell ([CL:0002242](https://www.ebi.ac.uk/ols/ontologies/cl/terms?iri=http%3A%2F%2Fpurl.obolibrary.org%2Fobo%2FCL_0002242)), a parent of lymphocyte ([CL:0000542](https://www.ebi.ac.uk/ols/ontologies/cl/terms?iri=http%3A%2F%2Fpurl.obolibrary.org%2Fobo%2FCL_0000542)). In general, these results showed that the presence of high *SPI1* expression proves sufficient to identify a cell type as hematopoietic.

CLA also selected *SPI1* in two cell type pairs that did not involve cell types from the hematopoietic cell lineage. The first pair comprises squamous epithelial cell ([CL:0000076](https://www.ebi.ac.uk/ols/ontologies/cl/terms?iri=http%3A%2F%2Fpurl.obolibrary.org%2Fobo%2FCL_0000076)) against multi fate stem cell ([CL:0000048](https://www.ebi.ac.uk/ols/ontologies/cl/terms?iri=http%3A%2F%2Fpurl.obolibrary.org%2Fobo%2FCL_0000048)). The second pair comprises cell types that come from the same two lineages as the first pair: epithelial cell ([CL:0000066](https://www.ebi.ac.uk/ols/ontologies/cl/terms?iri=http%3A%2F%2Fpurl.obolibrary.org%2Fobo%2FCL_0000066); parent of squamous epithelial cell) against mesenchymal cell ([CL:0008019](https://www.ebi.ac.uk/ols/ontologies/cl/terms?iri=http%3A%2F%2Fpurl.obolibrary.org%2Fobo%2FCL_0008019); child of multi fate stem cell). This suggests that *SPI1* expression can discriminate these two lineages.

Higher expression of *SPI1* in monocytes ([CL:0000576](https://www.ebi.ac.uk/ols/ontologies/cl/terms?iri=http%3A%2F%2Fpurl.obolibrary.org%2Fobo%2FCL_0000576)) discriminated against 51 of the 55 cell types compared. CLA did not use *SPI1* to discriminate monocytes against 4 other cell types: CD8-positive, alpha-beta T cell ([CL:0000625](https://www.ebi.ac.uk/ols/ontologies/cl/terms?iri=http%3A%2F%2Fpurl.obolibrary.org%2Fobo%2FCL_0000625)); blood cell ([CL:0000081](https://www.ebi.ac.uk/ols/ontologies/cl/terms?iri=http%3A%2F%2Fpurl.obolibrary.org%2Fobo%2FCL_0000081)); granulocyte ([CL:0000094](https://www.ebi.ac.uk/ols/ontologies/cl/terms?iri=http%3A%2F%2Fpurl.obolibrary.org%2Fobo%2FCL_0000094)); and B cell ([CL:0000236](https://www.ebi.ac.uk/ols/ontologies/cl/terms?iri=http%3A%2F%2Fpurl.obolibrary.org%2Fobo%2FCL_0000236)). These cell types be-long to the hematopoietic cell lineage (Figure 5), and we did not expect *SPI1* expression to discriminate monocytes against other hematopoietic cell types. As expected, *SPI1* had higher and discriminative expression exclusively in the hematopoietic lineage (Figure 5).

**Figure 5:**
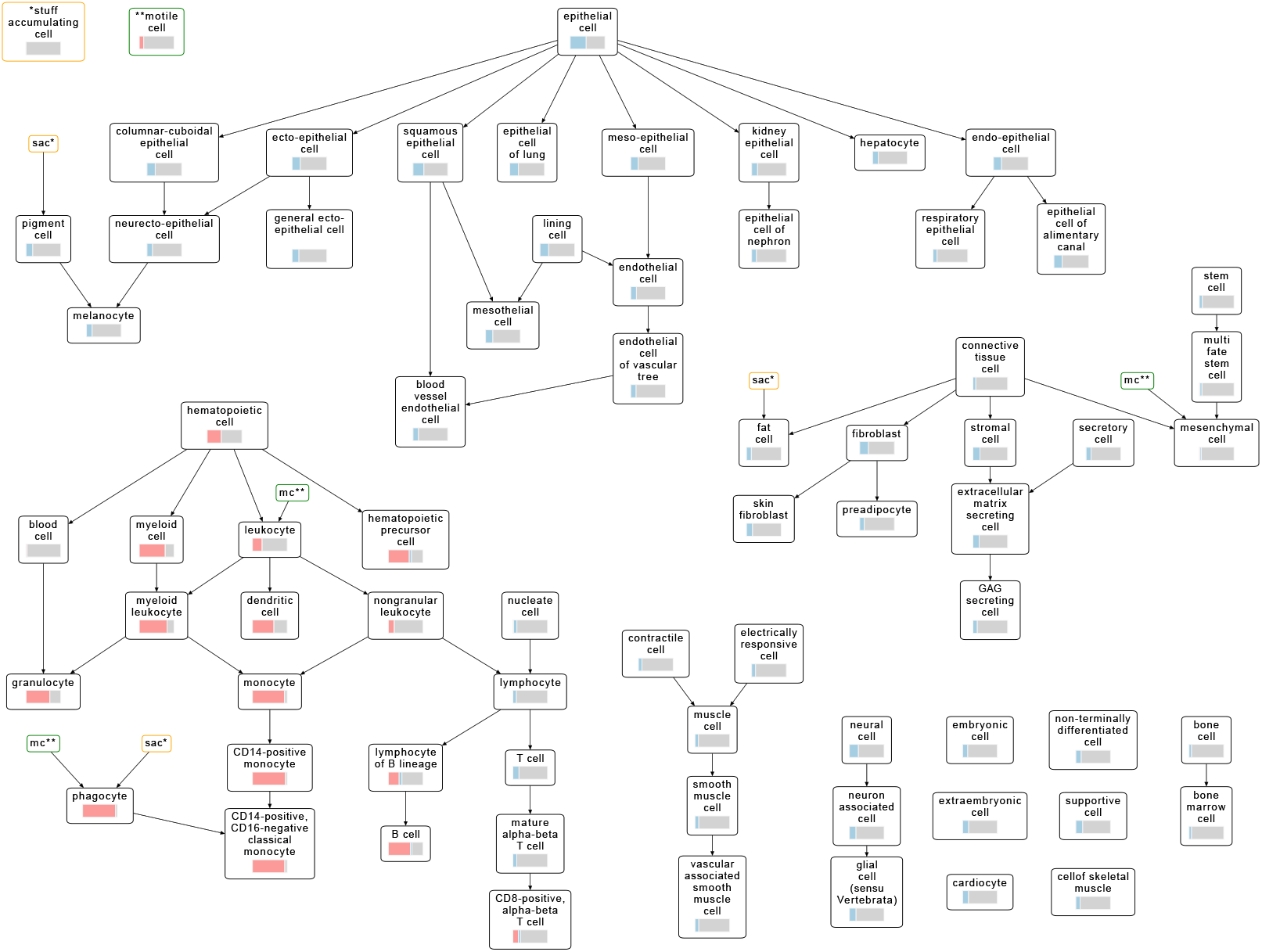
Lineage scores of the *SPI1* gene mapped to the Cell Ontology with a hierarchical layout. Nodes: cell types; edges: *is_a* relationship, with the arrow pointing to the descendant. Bar color: fraction of cell type comparisons where *SPI1* discriminates cell types and has higher expression in current cell type (red), higher expression in comparison cell type (blue), or no discriminative power (grey). The nodes for stuff accumulating cells (“sac*”, yellow) and motile cells (“mc**”, green) have descendants in multiple lineages. We duplicated these nodes to simplify the layout and show the lineage scores separately (top left).

Additionally, *SPI1* had low and non-discriminative expression in mature alpha-beta T cells and their ancestors, as described previously^18^. High expression in hematopoietic cells corresponds with known *SPI1* functions of positive regulation of monocytes, B cells, and dendritic cells^18,20^.

As *SPI1* had the highest lineage score (57/60) in CD14-positive monocytes ([CL:0001054](https://www.ebi.ac.uk/ols/ontologies/cl/terms?iri=http%3A%2F%2Fpurl.obolibrary.org%2Fobo%2FCL_0001054)), we also looked at the other transcription factors for this cell type. CLA identified the transcription factor *OLIG1* in more cell type pairs involving CD14-positive monocytes (59/60) than *SPI1*. This fits the known regulatory role of *OLIG1* transcription factor motifs in immune function and monocyte cells^21^.

### 2.4 CLA identifies cell-type–specific lncRNAs

The most common class of human genes, lncRNAs, have higher tissue specificity than protein-coding genes^22^. To identify cell-type-specific lncRNAs, we applied CLA to 4603 GENCODE lncRNA genes in FANTOM5 CAGE expression data for 71 cell types. We found 95 lncRNAs with CAGE expression that discriminates more than 80% of the comparisons for 15 cell types (Figure 6).

**Figure 6:**
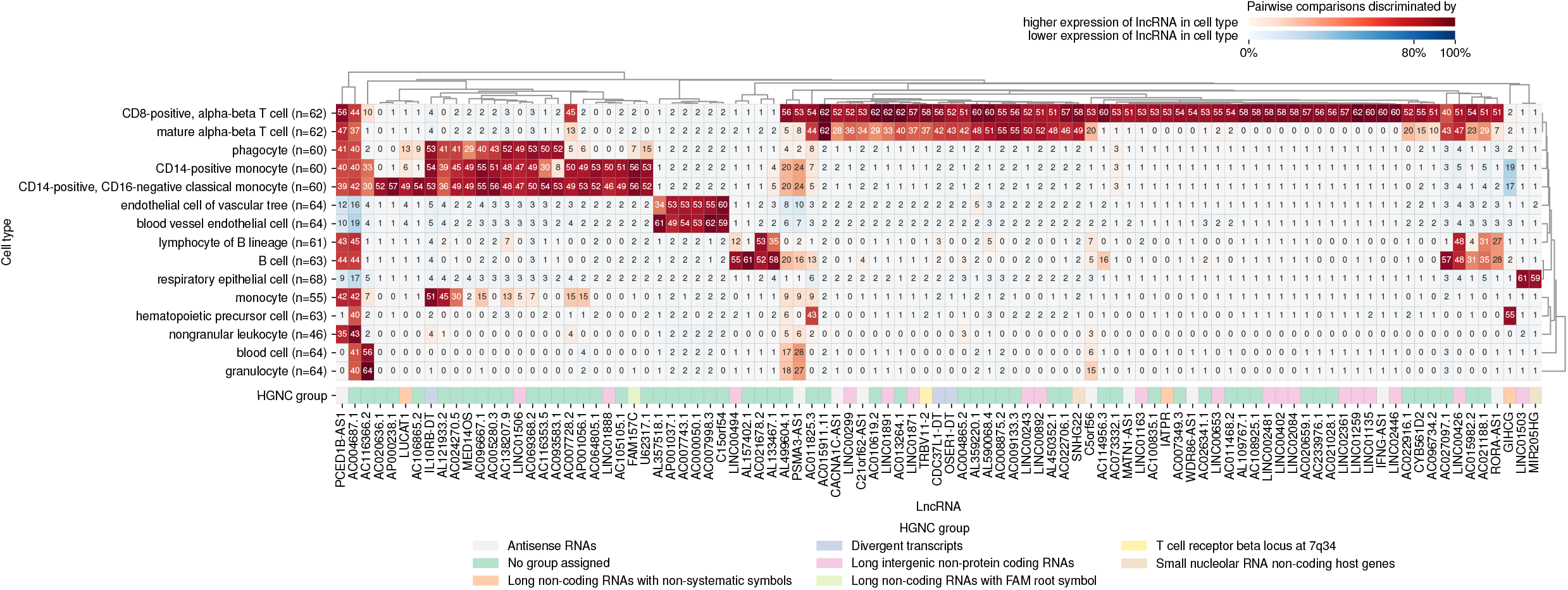
Heatmap of the 95 lncRNAs that best discriminate among 15 cell types. Both cell types (rows) and lncRNA genes (columns) ordered with hierarchical clustering using Euclidean distance. Cell color indicates that compared with other cell types, the lncRNA has higher (red), or lower (blue) expression in this cell type, or no discriminative power (white). Color intensity: percentage of comparisons discriminated by lncRNA per cell type. Number in cell: number of comparisons discriminated by this lncRNA per cell type. HGNC group: gene annotations by sequence similarity or function similarity from HUGO Gene Nomenclature Committee (HGNC).

Of the 95 cell-type–specific lncRNAs, 3 have a non-systematic HGNC^23^ gene symbol. These symbols denote genes with known function: *LUCAT1*, *IATPR*, and *GIHCG*.

The oncogene *LUCAT1*, known to promote multiple cancers^24–26^, has higher expression in whole blood^27^. It also has discriminative and higher expression in 49 pairwise comparisons (81%) with CD14-positive, CD16-negative classical monocytes ([CL:0002057](https://www.ebi.ac.uk/ols/ontologies/cl/terms?iri=http%3A%2F%2Fpurl.obolibrary.org%2Fobo%2FCL_0002057)).

Also regarded as an oncogene, *IATPR* has a role in breast cancer progression^28^ and non-small cell lung cancer progression^29^. CLA found *IATPR* with a discriminative higher expression in 53 comparisons (95%) with CD8-positive, alpha-beta T cell ([CL:0000625](https://www.ebi.ac.uk/ols/ontologies/cl/terms?iri=http%3A%2F%2Fpurl.obolibrary.org%2Fobo%2FCL_0000625)).

A third oncogene, *GIHCG*, with known involvement in hepatocellular carcinoma and squamous cell carcinoma^30,31^, had discriminative and higher expression in 55 pairwise comparisons (87%) of hematopoietic precursor cells ([CL:0008001](https://www.ebi.ac.uk/ols/ontologies/cl/terms?iri=http%3A%2F%2Fpurl.obolibrary.org%2Fobo%2FCL_0008001)).

Of the 95 lncRNAs that CLA selected, 57 had no HGNC group assigned. The cell-type specificity established by CLA along with the relative lack of information on the biological roles of these 57 lncRNAs make them good candidates for further investigation.

### 2.5 CLA identifies alternative promoters of cell surface markers

Cell surface proteins play an important role in cell communication and act as cell markers in flow cytometry^32^. We used CLA to identify cell-type–specific cell surface markers of immune cells, also known as CD genes^33^. To quantify promoter level expression we used FANTOM5 CAGE data, which quantify promoter expression at a higher resolution than RNA-seq^34^.

We used the FANTOM5 naming convention to distinguish alternative promoters^35^. This convention identifies each promoter by its rank of CAGE expression compared to other promoters from the same gene, followed by gene symbol. For example, *p1@CD53* refers to the most highly expressed promoter of the *CD53* gene.

As expected, the known B-cell biomarkers^36^ *CD19* and *TNFRSF17* (CD269)^37^ discriminated B cell ([CL:0000236](https://www.ebi.ac.uk/ols/ontologies/cl/terms?iri=http%3A%2F%2Fpurl.obolibrary.org%2Fobo%2FCL_0000236)) and lymphocyte of B lineage ([CL:0000945](https://www.ebi.ac.uk/ols/ontologies/cl/terms?iri=http%3A%2F%2Fpurl.obolibrary.org%2Fobo%2FCL_0000945)). The *CD53* gene’s most expressed promoter, p1@*CD53*, only discriminated cell type pairs that involved cell types from the hematopoietic cell lineage. Other *CD53* promoters, however, discriminated more specific cell types within the hematopoietic lineage. The third-most-expressed promoter, p3@*CD53*, discriminated only between B cells and their parent, lymphocyte of B lineage. The fifth-most-expressed promoter, p5@*CD53*, exclusively discriminated CD14-positive, CD16-negative classical monocytes ([CL:0002057](https://www.ebi.ac.uk/ols/ontologies/cl/terms?iri=http%3A%2F%2Fpurl.obolibrary.org%2Fobo%2FCL_0002057)). These results suggest that CLA identifies alternative promoters specific to particular cell types.

CLA found that for *CD22*, known as a B cell regulator^38^, promoter p1 *CD22* had discriminative expression only for B cells and its parent, lymphocyte of B lineage. Similarly to *CD53*, the discriminative power of *CD22*’s alternative promoter p13@*CD22* decreased in B cells and increased for CD14-positive, CD16-negative classical monocytes, and phagocytes ([CL:0000234](https://www.ebi.ac.uk/ols/ontologies/cl/terms?iri=http%3A%2F%2Fpurl.obolibrary.org%2Fobo%2FCL_0000234)).

### 2.6 Comparison of CLA against differential gene expression

We examined the utility of CLA for identifying biomarkers, as compared to differential expression analysis. Tools to perform differential expression analysis, such as DESeq26 or edgeR^7^, often return a long list of differentially expressed genes. We hypothesized that CLA’s regularized random forest approach might yield a smaller set of important genes with differential expression.

We compared the results of CLA with DESeq2 CAGE data for 1639 transcription factors in 29 samples associated with columnar-cuboidal epithelial cells ([CL:0002066](https://www.ebi.ac.uk/ols/ontologies/cl/terms?iri=http%3A%2F%2Fpurl.obolibrary.org%2Fobo%2FCL_0002066)) and 23 samples associated with skin fibroblasts ([CL:0002620](https://www.ebi.ac.uk/ols/ontologies/cl/terms?iri=http%3A%2F%2Fpurl.obolibrary.org%2Fobo%2FCL_0002620)). We chose these two cell types for two reasons. First, these two cell types have a relatively balanced number of samples associated. Second, these two cell types each inherit samples from a variety of leaf cell types (columnar-cuboidal epithelial cells: 10 leaf cell types; skin fibroblasts: 8 leaf cell types).

While DESeq2 identified 551 differentially expressed transcription factors, CLA identified 24 transcription factors. The 24 transcription factors selected by CLA did not contain transcription factors deemed equivalent (subsubsection 3.3.1). CLA identified 6 genes that occur in the top 10 DESeq2 results sorted by adjusted p-values (Figure 7a). The most differentially expressed gene using DESeq2, *PAX6*, had no expression in more than half of the samples of each group. Because of this, CLA selected *PAX6* only once out of 10 random forests. CLA always selected, however, significantly differentially expressed genes with a higher number of nonzero expression values, such as *MAF* or *RFX8*.

**Figure 7:**
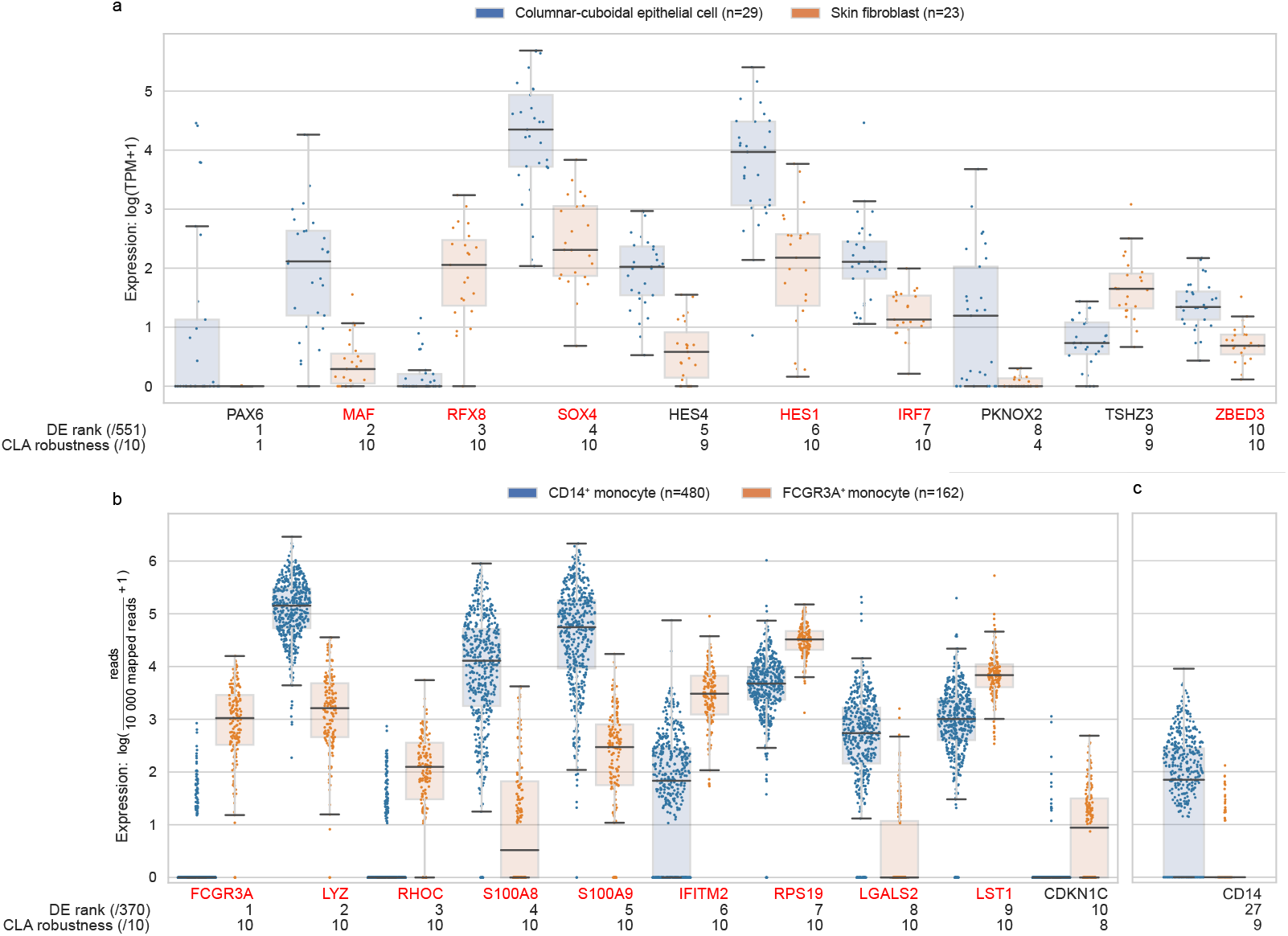
Expression of differentially expressed genes. Boxplot grey box: interquartile range (IQR). Boxplot whisker: most extreme value within quartile ±1.5 IQR. Sinaplot^39^: each dot represents a single cell. DE rank: rank of DESeq2 or Seurat adjusted p-value of significantly differentially expressed genes. Gene symbol color indicates CLA robustness score <10 (black) or exactly 10 (red). **(a)** Top 10 out of 551 significantly differentially expressed transcription factors reported by DESeq2 from 29 bulk CAGE columnar-cuboidal epithelial samples and 23 bulk CAGE skin fibroblast samples. **(b)** Top 10 out of 370 significantly differentially expressed genes reported by Seurat from single-cell RNA-seq data between 480 *CD14*^+^ monocyte samples and 162 *FCGR3A*^+^ monocyte samples. **(c)** *CD14*, the main driver of *CD14*^+^ monocyte samples, outside the top 10 differentially expressed genes reported by Seurat.

These results show that CLA identified fewer genes than differential expression methods. As CLA excluded the genes that often had zero expression in both cell types, it will prove more useful in many scenarios.

### 2.7 Identifying markers from single-cell data

After demonstrating CLA’s ability to identifying cell-type–specific in bulk primary cell and cell line data, we also tested it on single-cell RNA-seq data. We applied CLA to 10X Genomics single-cell RNA-seq data on 542 peripheral blood mononuclear cells (PBMCs) from a healthy donor. We repeated a scenario in the Seurat8 tutorial (https://satijalab.org/seurat/v3.2/pbmc3k_tutorial.html), finding genes that discriminate two clusters: CD14^+^ mono-cytes (n = 480) and FCGR3A^+^ monocytes (n = 162).

CLA identified 13 genes that discriminated between CD14^+^ monocytes and FCGR3A^+^ monocytes with robustness of 10/10. We compared the same cell types using Seurat^8^, which uses a non-parametric Wilcoxon rank sum test to identify differential expression in single-cell RNA-seq data. Seurat identified 370 significantly deferentially expressed genes. CLA’s selected genes included Seurat’s top 9 differentially expressed genes, as sorted by adjusted p-value. CLA’s results included the main driver of the FCGRA^+^ monocyte cluster *FCGR3A* (Figure 7b). The main driver of the CD14^+^ cluster, *CD14*, however, had a robustness score of 9/10, and had a lower corrected p-value from the results of differential expression using Seurat (Figure 7c).

The genes selected by CLA with robustness of 10/10 all have nonzero expression in more than 75% of the cells in at least 1 cluster. Alternatively, the genes selected with a robustness score lower than 10/10, such as *CD14*, had zero expression in more than 25% of the cells in both clusters. These results showed that CLA identifies cluster-specific genes not driven by zeros, unlike Seurat. Seurat also selected 370 genes, far more than the 13 genes selected by CLA.

## 3 Methods

### 3.1 Gene expression data

#### 3.1.1 Transcription factor promoters

We downloaded Ensembl^40^ gene identifiers for 1639 known and likely human transcription factors from the Human Transcription Factors database^41^ (https://humantfs.ccbr.utoronto.ca/download/v_1.01/TFs_Ensembl_v_1.01.txt). We identified genes with matching identifiers from the GENCODE^42^ v32 comprehensive gene set (ftp://ftp.ebi.ac.uk/pub/databases/gencode/Gencode_human/release_32/gencode.v32.annotation.gtf.gz), selecting rows with feature type gene using pybedtools^43^ v0.7.10. Then we used pybedtools to identify the first base of each gene, which we considered as the location of the promoter. With this approach, we associated each transcription factor with its gene’s most upstream promoter.

#### 3.1.2 lncRNA promoters

We identified genes with biotype lncRNA from the GENCODE^42^ v32 comprehensive gene set (ftp://ftp.ebi.ac.uk/pub/databases/gencode/Gencode_human/release_32/gencode.v32.annotation.gtf.gz), selecting rows with feature type gene using pybedtools^43^ v0.7.10. Then we used pybedtools to identify the first base of each gene, which we considered as the location of the promoter. With this approach, we associated each lncRNA with its gene’s most upstream one promoter.

#### 3.1.3 Cluster of differentiation (CD) promoters

We downloaded from the HGNC resource^23^ a list of 394 CD genes (HGNC group 471) in January 2020. We identified transcripts with matching identifiers from the GENCODE^42^ v32 comprehensive gene set (ftp://ftp.ebi.ac.uk/pub/databases/gencode/Gencode_human/release_32/gencode.v32.annotation.gtf.gz), selecting rows with feature type transcript using pybedtools^43^ v0.7.10. With BEDTools^44^ v2.27.1, we merged the CD promoters within 1000 bp using the command `bedtools merge -d 1000 -sorted`. This ensured that alternative promoters had at least 1000 bp distance between each other. We merged FANTOM5 CAGE peaks within 500 bp using the command `bedtools merge -d 500 -sorted`. With this approach, CD genes may have multiple promoters.

#### 3.1.4 Assignment of expression values to promoters

To assign an expression value to each promoter for each transcription factor, lncRNA, and CD, we extended their genomic coordinates by ±500 bp using pybedtools `slop`. Then, we intersected the promoters with FANTOM5 CAGE peaks (https://fantom.gsc.riken.jp/5/datafiles/reprocessed/hg38_latest/extra/CAGE_peaks_expression/hg38_fair+new_CAGE_peaks_phase1and2_tpm.osc.txt.gz) using the command `bedtools intersect -sorted - wo`. When multiple peaks overlapped a single promoter, we selected only the peak with the highest expression. This approach assigned promoter expression values for multiple promoters of the CD genes. For the transcription factor and lncRNA datasets, however, it assigned one expression value per gene.

### 3.2 Sample to cell type association

We associated 340 Cell Ontology cell types to 751 FANTOM5 samples using the rule that each parent cell type inherits all the samples from its descendants accessible via *is_a* relationships. We merged cell types associated with identical sets of samples. From the merged cell types, we identified the one individual cell type closest to the common ancestor in the Cell Ontology graph. We used this cell type’s name for the merged cell type. In case of tie, we use the name of first cell type in alphabetical order. To focus on major cell types, we removed Cell Ontology terms associated with fewer than 10 samples.

In total, we generated a list of samples for 71 distinct cell types. Each cell type had between 11 and 270 associated samples (mean: 45.9; median: 24).

### 3.3 Transcriptional feature selection with random forest

We used regularized random forest9 to identify a minimum set of transcriptional features that discriminate samples between two cell types.

#### 3.3.1 Identifying equivalently discriminative transcriptional features

We excluded transcriptional features with zero variance across all samples. To identify other features equivalent for discriminating between two cell types, we used the following process on the samples associated with either of the cell types:

1. Repeat 3 times:

a. Shuffle the list of samples, without replacement.
b. Encode the rank of the gene expression values for each transcriptional feature across the samples as a binary string:

i. Construct a single vector of gene expression values for that transcriptional feature across the samples for both cell types.
ii. Sort the vector in ascending order.
iii. Create a string of binary values from the sorted gene expression values, where the binary value indicates to which of the two cell types the sample corresponds.
c. Compute Hamming distance^45^ between every pair of binary strings.
2. Tag as equivalent those transcriptional features with Hamming distance of 0 in all 3 repetitions.
3. Consider each clique of mutually equivalent transcriptional features as an equivalence class.
4. Randomly select 1 feature to represent each equivalence class. Exclude the class’s other transcriptional features from the gene expression data.

The shuffling step in each iteration accounted for the arbitrary ordering of the sorted gene expression vectors in case of ties. Requiring a zero Hamming distance between two genes in 3 different random shufflings increased robustness in calling transcriptional features equivalent.

#### 3.3.2 Stratified bootstrap sampling

We used a stratified bootstrap sampling approach to reduce the bias in random forest construction that arises from class imbalance^17,46^. This approach guaranteed that each decision tree in each random forest started with an equal number of samples from each cell type—specifically the number of samples s from the cell type with the fewest samples. Specifically, we randomly selected, with replacement, s samples from each cell type. This means that each decision tree in each random forest might start with a slightly different collection of samples, possibly with some repetition of samples.

#### 3.3.3 Random forest out-of-bag score accuracy

We used out-of-bag samples from the bootstrap sampling step to provide an unbiased estimate of a random forest classifier, the out-of-bag score^12^. After growing a forest of decision trees, we classified each out-of-bag sample using the trees that grown without using the sample. If most trees correctly predicted the sample’s class, we considered the out-of-bag sample correctly classified. The proportion of out-of-bag samples that this procedure correctly classified is the out-of-bag score.

#### 3.3.4 Regularized random forest

The random forests consist of decision trees made of nodes (decision points) and edges (two branches from each decision point) that classify samples using *m* input transcriptional features. When constructing a decision tree from *2s* bootstrap-selected samples, we split nodes in a way that minimized the *Gini impurity*^47^, how often the decision tree would misclassify one of those training samples. In other words, we sought to find transcriptional features and expression thresholds that maximized the *Gini gain*, defined as the decrease in Gini impurity. Beginning with a root node containing all bootstrap selected input samples, we executed the following procedure on each node in a growing decision tree:

1. If all the samples in the node come from the same cell type, we consider it a leaf node and continue to the next unprocessed node.
2. We constructed a subset of the *m* input transcriptional features, consisting of a union of:

- 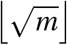 transcriptional features, randomly selected, without replacement
- all transcriptional features previously used to split nodes in this tree or any previously constructed trees
3. We split the node using the transcriptional feature in the subset that could maximize the Gini gain.

Only splitting nodes using transcriptional features that can provide larger gain than any previously used features provides a regularization capacity that reduces the total number of features used by the constructed random forest. Using a more compact set of features can reduce overfitting. We then reported these features along with the previously excluded features in their equivalence classes (subsubsection 3.3.1) as if CLA selected them. After this step, we did not distinguish the previously excluded features from other features.

#### 3.3.5 Robustness scores

Since random forest construction always begins with a random subset of transcriptional features, we might select features with low discriminative power by chance^48^. To identify and remove the selected transcriptional features with the lowest discriminative power, we defined a robustness score that indicated how many times we selected the transcriptional features in constructing 10 random forests, each forest itself made of many decision trees (Figure 3). We also reported a robustness score for all the transcriptional feature in equivalence classes.

### 3.4 Peripheral blood mononuclear cells (PBMCs)

We obtained exactly 2700 PBMC single-cell RNA-seq samples from 10X Genomics (https://cf.10xgenomics.com/samples/cell-exp/1.1.0/pbmc3k/pbmc3k_filtered_gene_bc_matrices.tar.gz). We processed the data using Seurat8 v4 and following the Seurat tutorial (https://satijalab.org/seurat/v3.2/pbmc3k_tutorial.html).

To remove multiplets, we discarded cells with more than 2500 genes detected. To remove low-quality cells, we discarded cells with fewer than 200 genes detected. To remove cells exhibiting mitochondrial contamination, we used the Seurat function `PercentageFeatureSet()` to discard cells with more than 5% of read counts overlapping with mitochondrial genes.

To normalize the data, we used the Seurat function `NormalizeData(normalizationmethod= ‘LogNormalized’, scale.factor=10000)`. This divided each gene read count by the number of reads in each cell, and multiplied the relative counts by 10 000.

### 3.5 Differential expression analysis

We used the Seurat function `FindMarkers()` with default parameters to identify the genes differently expressed between CD14^+^ monocytes and FCGR3A^+^ monocytes. We used the normalized PBMC data as input to Seurat (subsection 3.4).

We used DESeq2 v1.30 with default parameters to identify the transcription factors differentially expressed between columnar-cuboidal epithelial cells and skin fibroblasts. We used FANTOM5 read counts (https://fantom.gsc.riken.jp/5/datafiles/reprocessed/hg38_latest/extra/CAGE_peaks_expression/hg38_fair+new_CAGE_peaks_phase1and2_tpm.osc.txt.gz) to generate transcription factor promoter count data (subsubsection 3.1.4). We used the transcription factor promoter count data as input to DESeq2.

## 4 Discussion

We describe a simple, yet efficient, method to identify markers that distinguish between cell type pairs. Here, we used bulk CAGE and single-cell RNA-seq gene and promoter expression data, but one can easily apply our approach to other data containing genomic feature observations across large collections of samples.

While common differential expression tools selected more than 300 features on both single-cell RNA-seq and bulk CAGE datasets, CLA selected at most 24 features. Transcriptional features considered significantly differentially expressed by differential expression tools had median expression values of 0 in the two compared cell types, including the top transcriptional features (Figure 7). When features most often have expression values of 0 in both two cell types, they have decreased utility as distinctive markers for those cell types, to say the least. CLA, however, selected the top differentially expressed features without including zero-inflated features. In this way, CLA seems better at selecting cell-type–specific features. By selecting fewer and better cell-type–specific features, CLA simplifies analyses comparing multiple cell types. This makes CLA suitable for the analysis of large compendia of data describing many cell types.

Lacking comprehensive ground truth data on marker genes for cell types prevented hyperparameter tuning. Our method has one important hyperparameter, the number of trees used in each random forest. We chose to use 500 trees for every cell type pairs because this resulted in more comparisons with higher accuracy, as estimated using out-of-bag samples (Figure 2).

Using only the highest threshold (10/10) of robustness may eliminate important transcriptional features from selection by CLA. The regularization layer prevents the selection of transcriptional features less important than previously selected transcriptional features. Thus, by chance, one random forest instance may start using an important transcriptional feature in the first decision point and keep reusing it.

Using CLA, we automated the identification of cell-type–specific transcriptional features by comparing all possible cell type pairs from the FF sample ontology. Other studies that have used the Cell Ontology to identify cell-type–specific genes^49^ have required setting arbitrary thresholds for considering a gene as specific or not^50^ or have only considered increases in gene expression in a cell type of interest^51^. Our approach allows one to easily identify cell-type–specific transcriptional features from thousands of samples.

## 5 Acknowledgments

We thank Timo Lassmann (Telethon Kids Institute, University of Western Australia, 0000-0002-0138-2691) for helpful discussions throughout the project. This work was supported by the Natural Sciences and Engineering Research Council of Canada (RGPIN-2015-03948 to M.M.H.). We thank Carl Virtanen (0000-0002-2174-846X) and Zhibin Lu (0000-0001-6281-1413) at the University Health Network High Performance Computing Center and Bioinformatics Core for technical assistance.

## 7 Author contributions

Conceptualization, M.M. and M.M.H.; Data curation, M.M.; Formal analysis, M.M.; Funding acquisition, M.M.H.; Investigation, M.M.;Methodology, M.M. and M.M.H.;Project administration, M.M., and M.M.H.;Resources, M.M.H.; Software, M.M.; Supervision, M.M.H.; Validation, M.M. and M.M.H.; Visualization, M.M. and J.H.; Writing — original draft, M.M.; Writing — review & editing, M.M. and M.M.H.

